# Extreme restructuring of *cis*-regulatory regions controlling a deeply conserved plant stem cell regulator

**DOI:** 10.1101/2023.12.20.572550

**Authors:** Danielle Ciren, Sophia Zebell, Zachary B. Lippman

## Abstract

A striking paradox is that genes with conserved protein sequence, function and expression pattern over deep time often exhibit extremely divergent *cis*-regulatory sequences. It remains unclear how such drastic *cis*-regulatory evolution across species allows preservation of gene function, and to what extent these differences influence how *cis-*regulatory variation arising within species impacts phenotypic change. Here, we investigated these questions using a plant stem cell regulator conserved in expression pattern and function over ∼125 million years. Using *in-vivo* genome editing in two distantly related models, *Arabidopsis thaliana* (Arabidopsis) and *Solanum lycopersicum* (tomato), we generated over 70 deletion alleles in the upstream and downstream regions of the stem cell repressor gene *CLAVATA3* (*CLV3*) and compared their individual and combined effects on a shared phenotype, the number of carpels that make fruits. We found that sequences upstream of tomato *CLV3* are highly sensitive to even small perturbations compared to its downstream region. In contrast, Arabidopsis *CLV3* function is tolerant to severe disruptions both upstream and downstream of the coding sequence. Combining upstream and downstream deletions also revealed a different regulatory outcome. Whereas phenotypic enhancement from adding downstream mutations was predominantly weak and additive in tomato, mutating both regions of Arabidopsis *CLV3* caused substantial and synergistic effects, demonstrating distinct distribution and redundancy of functional *cis*-regulatory sequences. Our results demonstrate remarkable malleability in *cis*-regulatory structural organization of a deeply conserved plant stem cell regulator and suggest that major reconfiguration of *cis*-regulatory sequence space is a common yet cryptic evolutionary force altering genotype-to-phenotype relationships from regulatory variation in conserved genes. Finally, our findings underscore the need for lineage-specific dissection of the spatial architecture of *cis*-regulation to effectively engineer trait variation from conserved productivity genes in crops.

**Author summary:** We investigated the evolution of *cis*-regulatory elements (CREs) and their interactions in the regulation of a plant stem cell regulator gene, *CLAVATA3 (CLV3)*, in Arabidopsis and tomato. Despite diverging ∼125 million years ago, the function and expression of *CLV3* is conserved in these species; however, *cis*-regulatory sequences upstream and downstream have drastically diverged, preventing identification of conserved non-coding sequences between them. We used CRISPR-Cas9 to engineer dozens of mutations within the *cis*-regulatory regions of Arabidopsis and tomato *CLV3.* In tomato, our results show that tomato *CLV3* function primarily relies on interactions among CREs in the 5’ non-coding region, unlike Arabidopsis *CLV3*, which depends on a more balanced distribution of functional CREs between the 5’ and 3’ regions. Therefore, despite a high degree of functional conservation, our study demonstrates divergent regulatory strategies between two distantly related *CLV3* orthologs, with substantial alterations in regulatory sequences, their spatial arrangement, and their relative effects on *CLV3* regulation. These results suggest that regulatory regions are not only extremely robust to mutagenesis, but also that the sequences underlying this robustness can be lineage-specific for conserved genes, due to the complex and often redundant interactions among CREs that ensure proper gene function amidst large-scale sequence turnover.

## Introduction

*Cis*-regulatory control of gene expression is essential for the function of genes and the phenotypes they govern. Expression control depends on *cis*-regulatory elements (CREs), non-coding sequences of DNA bound by transcription factors that determine when, where, and to what level genes are expressed throughout growth and development. CREs can occur in many sequence contexts relative to the gene they regulate, including upstream (5’) and downstream (3’), within the gene itself (in UTRs, introns, and exons), and at distal sites far away. Molecular assays for chromatin accessibility, histone modifications, and transcription factor binding in many model organisms have been used to identify hundreds of thousands of putative CREs (1–7). Numerous reporter studies have been used to predict the effect of CREs on expression, and more recently functional genomics studies leveraging massively-throughput assays have dissected *cis*-regulatory control of gene expression at scale (2,8–13). In contrast, much less is known about how perturbation of *cis*-regulatory sequence space impacts phenotypes in multi-cellular organisms, both within and between species. Empowered by genome editing, studies can now go beyond the limited number and diversity of natural *cis*-regulatory alleles to address previously intractable questions on the intricate organization and relationships of CREs underlying genotype-to-phenotype relationships.

Compared to the strong selective pressures on protein-coding sequences, *cis*-regulatory regions and their modularly organized and often highly redundant CREs are much more tolerant to sequence change, and thus evolve more rapidly (14). Additionally, transcription factor binding sites (TFBSs) are degenerate, and their organization in spacing, order, orientation, and number is often highly flexible (15). As multiple sequence compositions can produce similar regulatory outcomes, identifying conserved non-coding sequences (CNSs) via conventional alignment strategies is difficult (16). This problem is even more apparent over longer evolutionary time scales, as *cis*-regulatory divergence between orthologous genes often results in little to no sequence similarity (17,18). Importantly, however, while natural variation in expression and phenotypes among related genotypes are most often associated with *cis*-regulatory change, affected genes typically do not deviate substantially from their original expression patterns and phenotypic consequences are largely limited to the tissues and organs in which the genes function (19–21). Moreover, co-expression and gene knockout studies across widely divergent species have found that for many orthologous genes, expression programs and phenotypes controlled are broadly conserved (22–25). Thus, how such deeply conserved genes can tolerate extreme *cis*-regulatory change but still maintain shared functions over deep time is an open question.

A prominent hypothesis is that despite overall sequence divergence, *trans*-factor identity along with the relative positioning and functional interactions among CREs are constrained to preserve control of orthologous genes over wide evolutionary distances (18,26). Along with fundamental insights into *cis*-regulatory evolution, understanding such constraints, often termed as “grammar” (15), could accelerate efforts to transfer knowledge from model organisms to new species, especially for trait engineering. We addressed this question by taking advantage of *Arabidopsis thaliana* (Arabidopsis) and *Solanum lycopersicum* (tomato), two plant model systems separated by ∼125 million years of evolution that offer equally powerful tools in genome editing and high throughput phenotyping (27). We leveraged the *CLAVATA3* (*CLV3*) gene, which encodes a functionally conserved signaling peptide that represses stem cell proliferation in a deeply conserved negative feedback loop with the stem cell promoting transcription factor gene *WUSCHEL* (*WUS*) (28). Null mutations of *CLV3* in both systems cause the same stem cell over-proliferation phenotypes, most evident in an increase in the number of floral organs, including the carpels that form seed compartments in fruits known as locules (28). Using *in vivo* CRISPR-Cas9 genome editing of *CLV3* in both species, we identify CREs, resolve their organization, and dissect their interactions. This comparative approach allowed direct assessment of genotype-to-phenotype relationships in an evolutionary context, revealing the dynamics of *cis*-regulatory change of a conserved gene over deep time.

## Results

### The *cis*-regulatory sequences of *CLV3* in Arabidopsis and tomato are highly diverged

The signaling peptide CLV3 is a conserved 12 amino acid negative regulator of meristem proliferation in flowering plants (**Fig 1A**) (28,29). Loss of *CLV3* in both Arabidopsis and tomato results in overproliferation (fasciation) of shoot tissue and also organ number in flowers and fruits (**Fig 1B, C**). Consistent with its shared developmental role, both genes have similar expression domains in the shoot meristem, and both dodecapeptides are modified with an arabinose sugar chain and bind to orthologous receptor-like kinases to repress *WUS* (**Fig 1D, E**) (30–32). These similarities in expression, function, and mutant phenotypes made *CLV3* an ideal system to investigate *cis*-regulatory evolution of a conserved gene over long evolutionary time.

**Fig 1.**
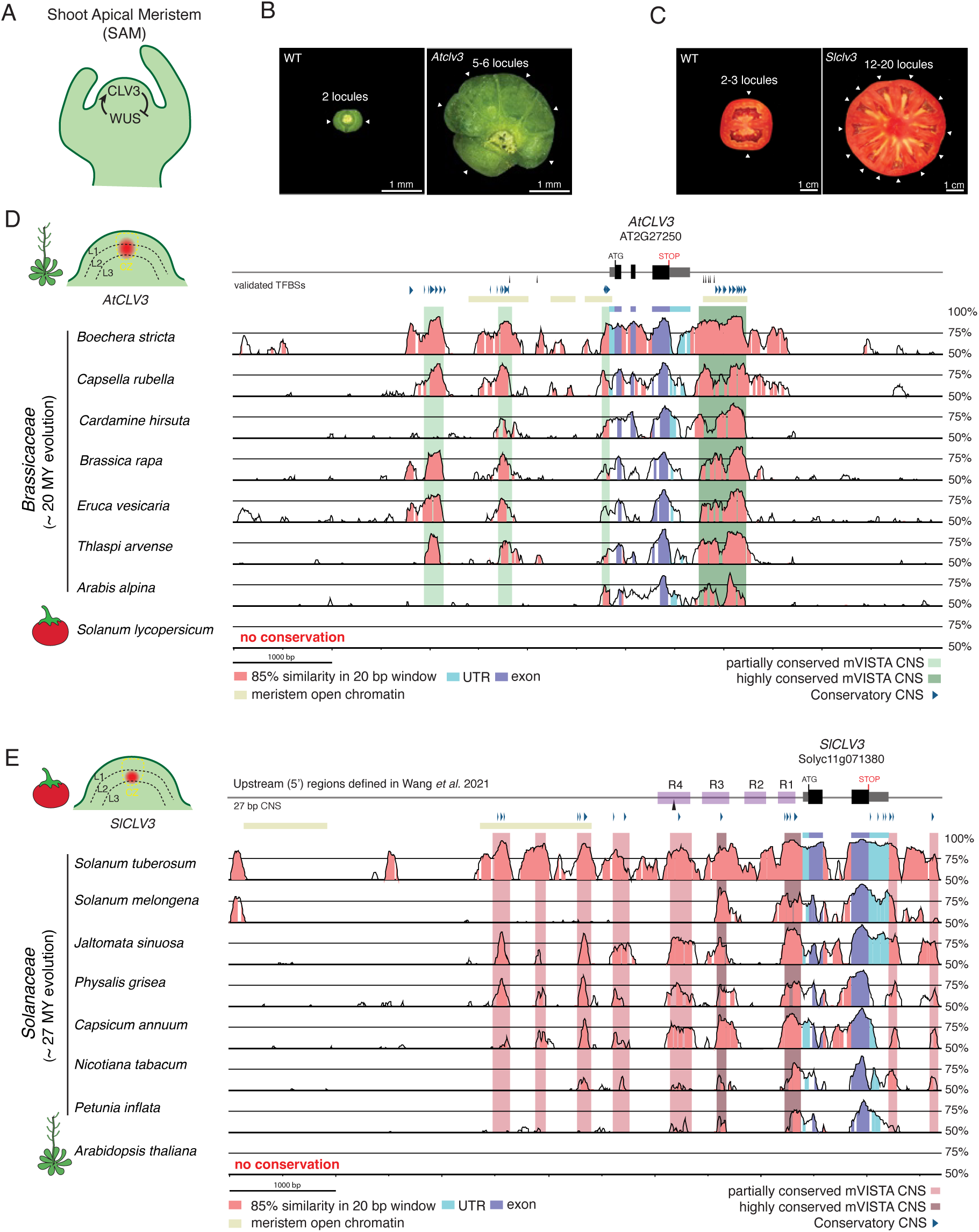
The function of CLV3 in Arabidopsis and tomato is conserved despite extreme divergence in *cis*-regulatory sequences. (A) A representative shoot apical meristem (SAM), demonstrating the conserved negative feedback loop between the signaling peptide CLV3 and the transcription factor WUS. CLV3 peptide indirectly inhibits WUS expression, while WUS promotes CLV3 expression. (B) Top-down view of Arabidopsis siliques from wild type (WT) and an *Atclv3* null mutant. White arrows, individual locules. Scale bars, 1 mm. (C) Transverse sections of tomato fruits from WT and a *Slclv3* null mutant. White arrows, individual locules. Scale bars, 1 cm. (D) *AtCLV3* gene model and surrounding regulatory regions upstream and downstream. mVISTA DNA sequence alignments of seven *CLV3* orthologs from Brassicaceae species, using the *AtCLV3* gene and its surrounding regulatory regions as the reference sequence*. SlCLV3* could not be aligned to *AtCLV3.* Sequences conserved in all species are denoted by a dark green bar, and sequences conserved in at least half of the species are denoted by a light green bar. A representative diagram of the Arabidopsis SAM is shown to the left, indicating the location of *CLV3* RNA expression relative to previously defined regions. L1, L2, and L3 layers are denoted by dotted black lines, the central zone (CZ) is outlined in yellow, and *CLV3* transcripts are represented in red. (E) *SlCLV3* gene model and surrounding regulatory regions upstream and downstream. mVISTA DNA sequence alignments of seven *CLV3* orthologs from Solanaceae species, using the *SlCLV3* gene and its surrounding regulatory regions as the reference sequence*. AtCLV3* could not be aligned to *SlCLV3.* Sequences conserved in all species are denoted by a dark red bar, and sequences conserved in at least half of species are denoted by a light red bar. Regions of the *SlCLV3* promoter previously defined are highlighted in purple (34). A representative diagram of the tomato SAM is shown to the left, indicating the location of *CLV3* RNA expression relative to previously defined regions. (D) – (E) Conservation was calculated as sequences with 85% similarity in 20 bp windows. The gene models are annotated with the location of previously validated TFBSs (black arrows), meristem open chromatin (yellow bars), and conserved non-coding sequences (CNSs) (blue arrows) defined by Conservatory (24,35,36). Light blue, conserved UTRs. Dark blue, conserved exons. Pink, conserved regions.

We first sought to identify conserved non-coding sequences (CNSs) by aligning the regulatory regions of Arabidopsis and tomato *CLV3* (denoted *AtCLV3* and *SlCLV3*) with several related species in their respective Brassicaceae and Solanaceae families using mVISTA (33). While we identified several regions of partially and highly conserved non-coding sequence across species within each family, there was no similarity when comparing the conserved regions between Arabidopsis and tomato (**Fig 1D, E**). Our independent *cis*-regulatory conservation analysis method, Conservatory, defined short CNSs within the *AtCLV3* or *SlCLV3* regulatory regions that broadly overlapped with mVISTA CNSs (24). Notably, Conservatory CNSs were only discovered at the family and order level in Arabidopsis, and the family level in tomato, with none shared between Arabidopsis and tomato. Interestingly, the Brassicaceae CNSs of *AtCLV3* were evenly distributed between the upstream and downstream surrounding sequence, whereas the Solanaceae CNSs were more abundant upstream of *SlCLV3* relative to downstream (**S1 Table**). We explored whether transcription factor motifs were shared between Brassicaceae and Solanaceae Conservatory CNSs through FIMO motif enrichment analysis. While this analysis revealed few overlapping motifs, they were distributed differently in each species, with the shared motifs found both upstream and downstream of *AtCLV3* compared to a bias for shared motifs upstream of *SlCLV3* (**S1 Table**). Thus, the *cis*-regulatory regions controlling Arabidopsis and tomato *CLV3* appear highly diverged, and the altered spatial distribution of family-wide conservation suggests that the grammar underlying *CLV3 cis-*regulation may have been re-organized.

### Neither the region upstream or downstream of *AtCLV3* is essential for its function

To understand how such divergent non-coding regions nevertheless support similar gene functions, we used genome editing to dissect the *cis*-regulatory functional organization of *SlCLV3* and *AtCLV3*. We previously used CRISPR-Cas9 multiplex mutagenesis to create an allelic series in the proximal 2 kb upstream of *SlCLV3*, which revealed several critical regulatory regions, including proximal and distal sequences that when deleted together phenocopy *Slclv3* null mutant effects on locule number (34,37). Using the same unbiased approach in Arabidopsis, we designed two 8-gRNA arrays to generate deletions within a 1.5 kb region upstream of the *AtCLV3* transcription start site (TSS), and also throughout the entire 3.8 kb region between the TSS and the upstream gene (**Fig 2A**). We generated 11 *AtCLV3^Reg5’^* alleles and assessed their effects by quantifying locule number in homozygous mutants. In striking contrast to our findings in tomato, all 11 alleles had little or no effect on locule number. For example, the most significant effect came from *AtCLV3^Reg5’-11^*, which despite losing nearly the entire 3.8 kb target region and all CNSs produced only three or four locules in half of its fruits, compared to an average of two locules in WT fruits and more than five in *Atclv3* null mutants (**Fig 2A**). Moreover, while substantial locule number increases emerged from deletions in the distal 5’ region of *SlCLV3*, the two *AtCLV3* alleles with deletions affecting large portions of the distal target regions showed no phenotypes (*AtCLV3^Reg5’-9,10^*). Finally, two proximal deletion alleles (*AtCLV3^Reg5’-3,7^*) exhibited only a slight increase in locule number, despite being within 100 bp of the TSS (**Fig 2A, B**). Thus, unlike in tomato, the *AtCLV3* 5’ region is mostly dispensable for maintaining regulation and function.

**Fig 2.**
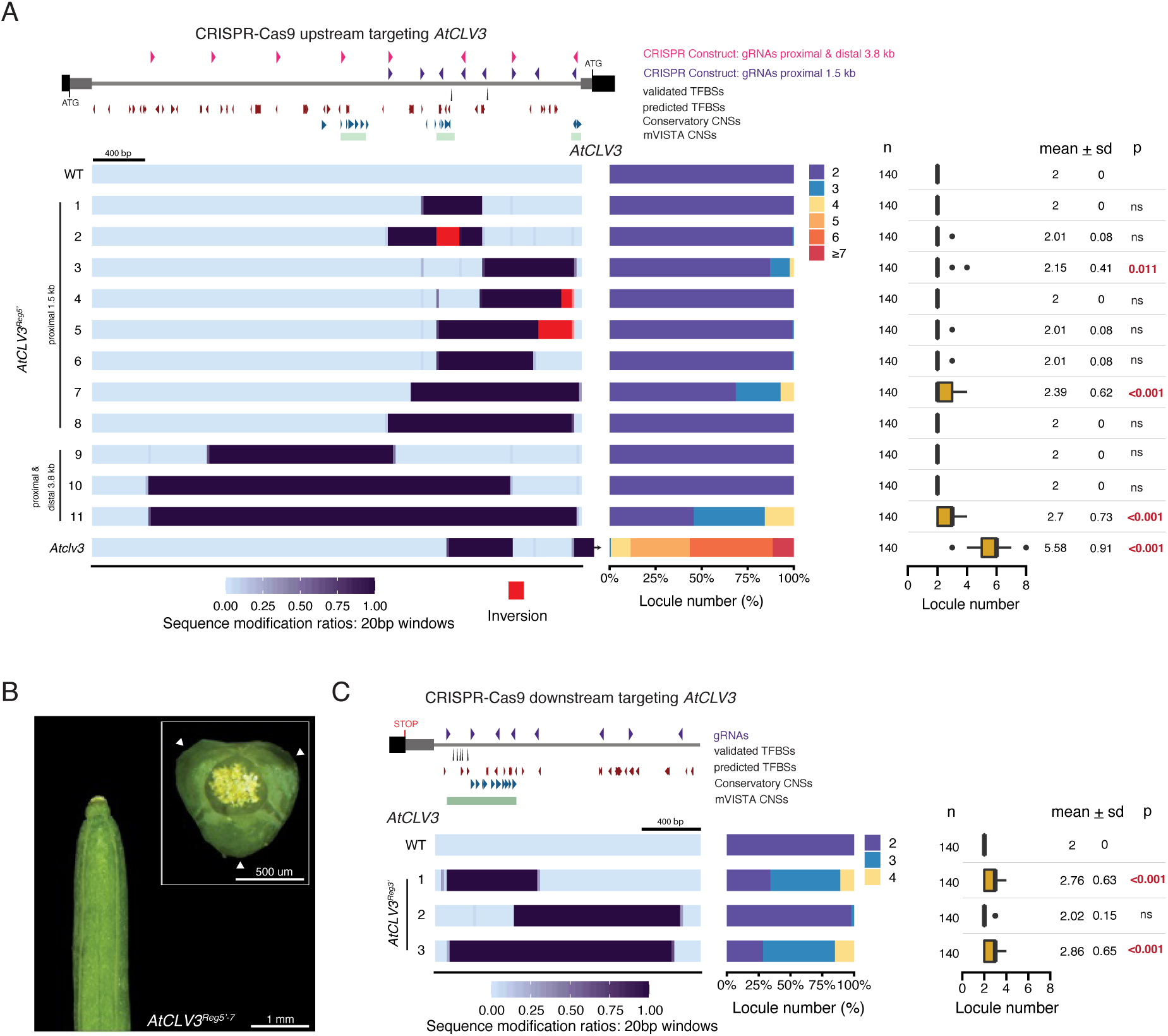
CRISPR-Cas9 mutagenesis of regions upstream or downstream of *AtCLV3* reveal that neither region alone is essential to its function. (A) Representation of the *AtCLV3* gene (right) and the 3.8 kb region up to the next upstream gene (left). The gRNA array spanning the 1.5 kb sequence proximal to the TSS of *AtCLV3* is shown as purple arrowheads, and the gRNA array targeting the entire 3.8 kb region between the TSS of *AtCLV3* and the next gene upstream is shown as pink arrowheads. A heatmap representation of the 11 CRISPR-Cas9 engineered 5’ *AtCLV3^Reg5’^*alleles including WT and *Atclv3* null mutant. (B) A silique with three locules from the 5’ allele *AtCLV3^Reg5’-7^*. A top-down view is shown in the inset. Scale bar, 1 mm. Inset scale bar, 500 um. (C) Representation of the *AtCLV3* gene (left), and the 1.6 kb region downstream. gRNAs are represented by purple arrowheads. Heatmap representation of the three engineered 3’ *AtCLV3^Reg3’^*alleles. (A), (C) The alleles have been encoded, such that perturbations to the region are represented as the degree of sequence modification relative to WT within 20 bp windows. Inversions are shown in red in the encoding. The location of validated TFBSs (black arrows), TFBSs predicted by FIMO (red arrowheads), and CNSs identified by Conservatory (blue arrowheads) and mVISTA (green bars) are shown. Locule number quantifications are represented by stacked bar plots and box plots. Box plots show the 25^th^, 50^th^ (median) and 75^th^ percentiles, with outliers as black points. Number of fruits sampled (n) is shown to the left, and mean and standard deviation (sd) are shown to the right. Two-sided Dunnett’s compare with control tests were performed to compare engineered alleles to WT, and the p-values (p) are included to the right. ns, not significant.

Since large sequence perturbations 5’ of *AtCLV3* produced weak phenotypes, and CNSs were distributed both 5’ and 3’ (**S1 Table**), we hypothesized that critical CREs may be present downstream. In support, previous transgenic reporter assays and complementation showed that *AtCLV3* requires 1.2 kb of the 3’ region in order to recapitulate endogenous expression (38). In addition, chromatin immunoprecipitation (ChIP) experiments confirmed WUS protein binds to canonical WUS TFBSs located both 5’ and 3’ of *AtCLV3,* and the transcription factor meristem regulator SHOOT MERISTEMLESS (STM) binds at a 5’ site (**Fig 2A, C**) (35,36,39). Five of the six WUS binding sites are clustered within a 116 bp region 3’ of *AtCLV3,* and previous work characterized a *cis-*regulatory module involving WUS cooperativity and concentration to modulate expression domain and level of *AtCLV3* (35).

By targeting 1.6 kb of the 3’ region proximal to the 3’UTR of *AtCLV3,* including the WUS binding sites, we isolated three large deletion alleles, one proximal, one distal, and one that nearly completely removed the targeted region (**Fig 2C**). Only the proximal and complete deletion alleles (*AtCLV3^Reg3’^*^-*1*^ and *AtCLV3^Reg3’^*^-3^) had weak phenotypes, and both resembled *AtCLV3^Reg5’^*^-*11*^, the largest 5’ deletion allele. Notably, both *AtCLV3^Reg3’-1^* and *AtCLV3^Reg3’-3^* lack the WUS binding sites and all 3’ CNSs. Altogether, these results demonstrate that while large portions of *AtCLV3* 5’ and 3’ regions are required to maintain complete wild type function, each region is largely dispensable.

### *AtCLV3* function depends on redundant CREs partitioned upstream and downstream

We previously showed that *SlCLV3* 5’ proximal and distal regions function in multiple complex functional relationships, including additive, redundant and synergistic interactions between *cis*-regulatory sequences, revealed by combining specific mutations (34). The absence of strong phenotypes from eliminating large portions of the 5’ or 3’ *cis*-regulatory regions of *AtCLV3* suggested there might be interactions between these regions. To test this, we took two parallel approaches to generate additional mutant alleles with perturbations in both 5’ and 3’ regions. We selected two large deletion alleles proximal to the TSS (*AtCLV3^Reg5’^*^-*8*^ and *AtCLV3^Reg5’^*^-*7*^) and transformed them with the 8-gRNA array previously used to generate 3’ mutations (**Fig 3A**). In a complementary approach, we transformed our largest 3’ deletion allele (*AtCLV3^Reg3’-3^*) with the 8-gRNA array previously used to generate 5’ mutations proximal to the TSS. From both experiments, we generated 28 new *cis*-regulatory alleles with various combinations of 5’ and 3’ mutations (**Fig 3B, C, D**). We then examined interaction effects between specific 5’ and 3’ mutations to determine if the enhancement in locule number from combined mutations was equal to the sum of the individual effects of 5’ and 3’ mutations (additive) or greater than the combined effects of each allele (synergistic or redundant).

**Fig 3.**
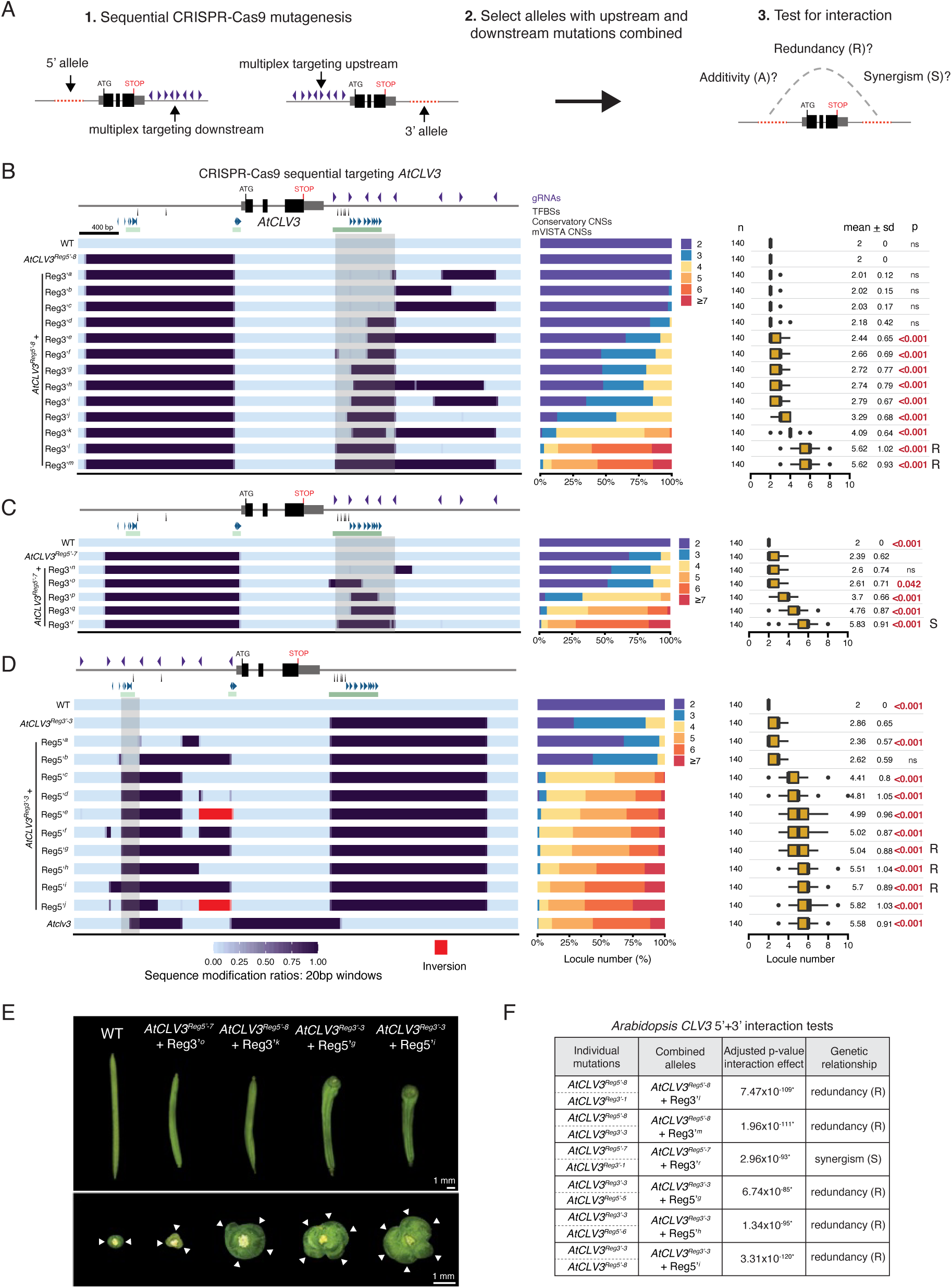
Combined mutagenesis of regions upstream and downstream of *AtCLV3* uncovers functional redundancy between these regions. (A) Schematic describing sequential CRISPR-Cas9 mutagenesis technique. Either a fixed 5’ allele was transformed with 3’-targeted gRNAs to induce new 3’ mutations, or a fixed 3’ allele was transformed with 5’-targeted gRNAs to induce new 5’ mutations. Transgenics were screened for new mutations, and alleles with both 5’ and 3’ mutations were selected. Genetic interaction tests were applied to explore the relationship between combined mutations in the 5’ and 3’ regions. (B) Heatmap representation of alleles generated from sequential CRISPR-Cas9 targeting with the *AtCLV3* 3’-gRNA array, in the background of the fixed 5’ mutant *AtCLV3^Reg5’-8^*, and their locule number quantifications. (C) Heatmap representation of alleles generated from sequential CRISPR-Cas9 targeting with the *AtCLV3* 3’-gRNA array, in the background of the fixed 5’ mutant *AtCLV3^Reg5’-7^*, and their locule number quantifications. (D) Heatmap representation of alleles generated from sequential CRISPR-Cas9 targeting with the *AtCLV3* proximal 1.5 kb 5’-gRNA array, in the background of the fixed 3’ mutant *AtCLV3^Reg3’-3^*, and their locule number quantifications. (E) Representative silique images from WT and several alleles generated through sequential CRISPR-Cas9 editing. A top-down view is shown below. White arrows, individual locules. Scale bars, 1 mm. (F) Interaction tests performed between combined 5’+3’ alleles and similar individual 5’ and 3’ mutants. p-values of the interaction effect were adjusted for multiple comparisons. (B) – (D) Locule number quantifications are represented by stacked bar plots and box plots. Box plots show the 25^th^, 50^th^ (median) and 75^th^ percentiles, with outliers as black points. Number of fruits sampled (n) is shown to the left, and mean and standard deviation (sd) are shown to the right. Grey boxes highlight identified regions of importance for regulation. R, redundant interaction type. S, synergistic interaction type. A, additive interaction type. Purple arrowheads, gRNAs. Black arrows, validated WUS and STM TFBSs. Blue arrowheads, Conservatory CNSs. Green bars, mVISTA CNSs. Two-sided Dunnett’s compare with control tests were performed to compare WT and sequentially edited alleles to (B) *AtCLV3^Reg5’-8^*, (C) *AtCLV3^Reg5’^*^-*7*^, or (D) *AtCLV3*^Reg3’-3^.

Unlike alleles disrupting the 5’ or 3’ regions alone, this series of 28 combinatorial alleles spanned the entire spectrum of variation for locule number, including a null-like phenotype. Notably, in addition to demonstrating strong synergistic interactions between upstream and downstream mutations, the breadth of this allelic series allowed us to identify specific subregions and their associated sequences that interact redundantly to control *AtCLV3*. For instance, new 3’ targeting in the background of the 5’ allele *AtCLV3^Reg5’-8^* revealed that deletion of the ∼600 bp region between gRNA-1 and gRNA-5 downstream of *AtCLV3* is sufficient to produce a null-like phenotype in this 5’ mutant background (*AtCLV3^Reg5’-8^* + Reg3’*^l^*) (**Fig 3B**). Notably, this region eliminates the 3’ cluster of WUS TFBSs and flanking sequence, whereas partial deletions of this region, as well as mutations distal to this region, only had weak or moderate effects on locule number in the 5’ mutant background. Importantly, the strong effect from removing the ∼600 bp region was also found by generating new 3’ mutations in the background of the 5’ allele *AtCLV3^Reg5’-7^* (*AtCLV3^Reg5’^*^-7^ + Reg3’*^r^*) (**Fig 3C**). While an allele with a smaller deletion within this 600 bp region that removed all five 3’ WUS binding sites (*AtCLV3^Reg5’-7^* + Reg3’*°*) caused a slight enhancement in locule number, a strong synergistic and null-like effect required also deleting the sequence adjacent to the 3’ WUS TFBSs that encompasses a cluster of CNSs (*AtCLV3^Reg5’^*^-*7*^ + Reg3’*^p^* and *AtCLV3^Reg5’-7^* + Reg3’*^q^*) (**Fig 3C**, **E**). To minimize allele-specific effects, we focused our statistical analysis of interaction between the 5’ and 3’ alleles on the combined alleles that were most similar to the corresponding deletion in the individual allelic series. In all cases, the combined effects of 5’+3’ mutations were non-additive, revealing a range of synergistic effects due to redundancy between 5’ and 3’ CREs of *AtCLV3* (**Fig 3F, S1 Fig**).

We were also able to further functionally dissect the *AtCLV3* 5’ region by sequential mutagenesis in the background of the 3’ allele *AtCLV3^Reg3’-3^* (**Fig 3D**). The majority of these alleles deleted the 5’ WUS binding site as well as multiple Conservatory CNSs and had moderate or strong phenotypes. One combined allele (*AtCLV3^Reg3’-3^* + Reg5’*^b^*) deleted a large 5’ region but left the 5’ WUS site intact and perturbed fewer CNSs and had no effect. This suggests the increased importance of the 5’ WUS binding site and/or other conserved 5’ CREs in the absence of certain 3’ CREs. All combined alleles demonstrated redundant interactions between 5’ and 3’ mutations (**Fig 3F, S1 Fig**).

Altogether, these data show that alleles combining deletions both in 5’ and 3’ regions produce a range of *CLV3* loss-of-function phenotypes, demonstrating that *cis-*regulatory control of *AtCLV3* is partitioned between CREs upstream and downstream interacting synergistically to ensure wild type function. Our depth of allelic diversity allowed mapping of critical CREs to a ∼200 bp region upstream of *AtCLV3* and a ∼600 bp region downstream (highlighted in grey), notably overlapping with clusters of CNSs (**Fig 3B, C, D**). CREs within these regions contribute to *AtCLV3* function, however additional CREs upstream likely also influence *AtCLV3* regulation.

### The region downstream of *SlCLV3* contributes negligibly to its function

The partitioning and redundancy of critical 5’ and 3’ CREs controlling Arabidopsis *CLV3* contrasted with our previous findings in tomato, which suggested CRE function was restricted to the upstream region (34,37). In contrast to Arabidopsis, complete deletion of the 5’ targeted region of *SlCLV3,* which includes four CNS regions, produced a null-like phenotype on locule number (**Fig 4A, B**). Smaller deletions that overlapped with individual CNSs showed a continuum of less severe effects, with a general trend of distal deletions having greater consequences than proximal deletions (**Fig 4A, B**). However, combining smaller CNS deletions resulted in mostly weakly additive or redundant effects, suggesting broad dispersal of functional CREs across the 5’ region of *SlCLV3* (34).

**Fig 4.**
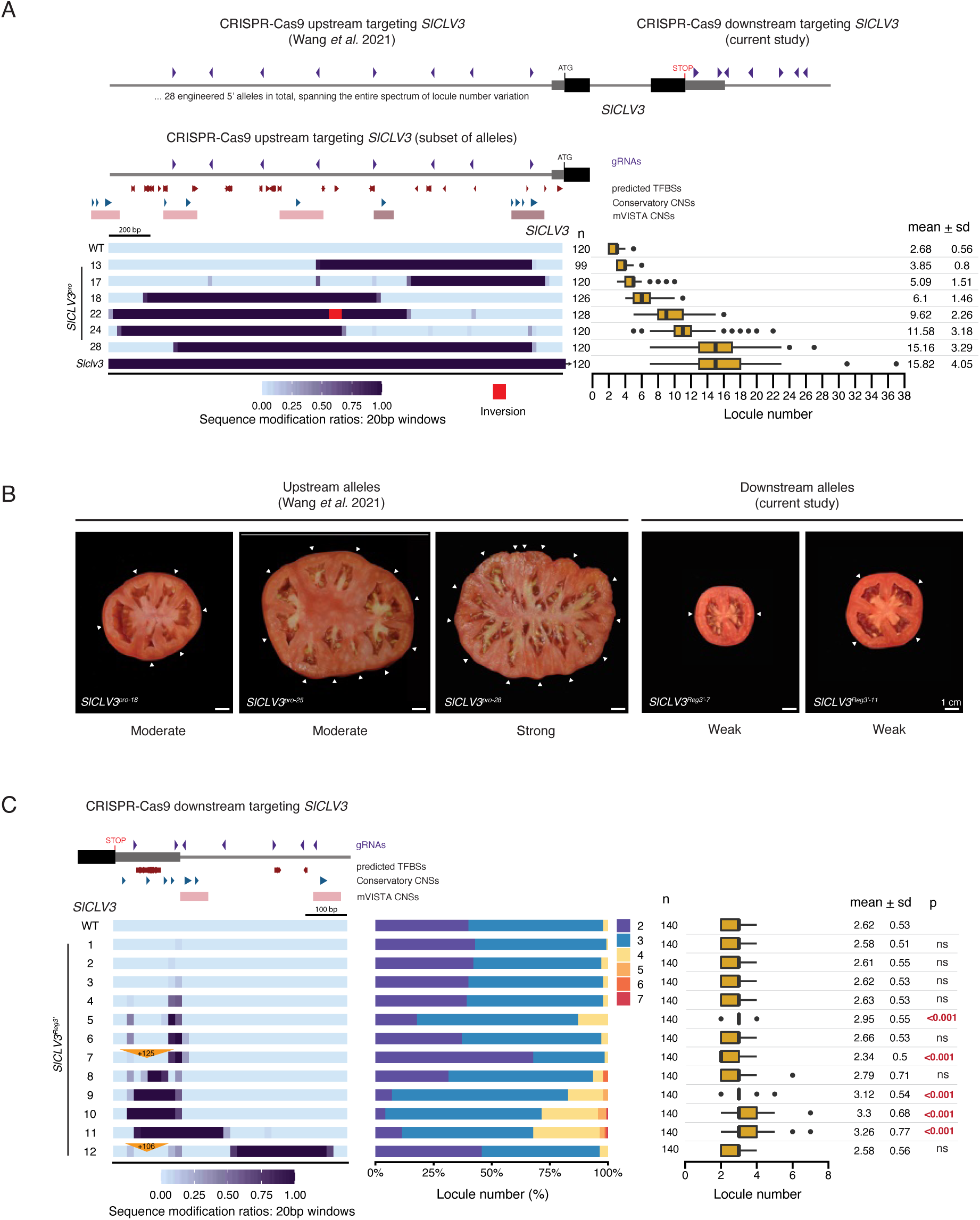
The region downstream of *SlCLV3* minimally contributes to gene function. (A) Schematic depicting the gRNA arrays (purple arrowheads) used to engineer mutations in the *SlCLV3* 5’ and 3’ non-coding regions using CRISPR-Cas9, in a past study and this study. The graph below reproduces a previous analysis of the 5’ non-coding region of *SlCLV3*, with a heatmap representation of a subset of the 28 alleles produced (34). Locule number quantifications are represented by box plots. (B) Representative images of tomatoes generated from 5’ or 3’ *SlCLV3* CRISPR-Cas9 mutagenesis. White arrows, individual locules. Scale bar, 1 cm. (C) Representation of the *SlCLV3* gene (left), and the gRNAs (purple arrowheads) used to engineer mutations in the region downstream. Heatmap representation of the 12 CRISPR-Cas9 engineered 3’ *SlCLV3^Reg3’^*alleles. Insertions are represented by orange triangles. Locule number quantifications are represented by stacked bar plots and box plots. A two-sided Dunnett’s compare with control test was performed to compare all 3’ engineered alleles to WT. (A), (C) Box plots show the 25^th^, 50^th^ (median) and 75^th^ percentiles, with outliers as black points. Number of fruits sampled (n) is shown to the left, and mean and standard deviation (sd) are shown to the right. Red arrows, predicted TF motifs from FIMO. Blue arrowheads, Conservatory CNSs. Red bars, mVISTA CNSs.

CNSs are also located downstream of *SlCLV3,* which could have *cis*-regulatory functions alone or interact with 5’ regions. In particular, a proximal region that overlaps with the 3’ UTR includes multiple CNSs and predicted TFBSs. We used multiplexed CRISPR-Cas9 mutagenesis to generate 12 3’ alleles with a range of sequence perturbations (**Fig 4C**). Similar to Arabidopsis, all alleles showed weak or no effects on locule number, but notably the three alleles with the most substantial phenotypes (*SlCLV3^Reg3’-9,10,11^*) affected multiple CNSs and predicted TFBSs. Interesting, *SlCLV3^Reg3’-7^* showed a greater proportion of two-locule fruits compared to WT, indicating a weak gain-of-function phenotype caused by a small deletion and a 125 bp duplication insertion that also duplicated several predicted TFBSs, which when lost cause a weak phenotype. Finally, despite removing half of the targeted region and the entire distal 3’ region including a CNS, allele *SlCLV3^Reg3’-12^* resembled WT. However, this allele also includes a similar duplication insertion to *SlCLV3^Reg3’-7^*, which could be suppressing a weak increase in locule number. Overall, these findings reveal a minor contribution of the 3’ *cis*-regulatory region in controlling *SlCLV3* function, in striking contrast to the organization of *AtCLV3 cis*-regulatory control.

### Upstream CREs have a dominant role in regulating *SlCLV3*, exhibiting weak interactions with downstream CREs

Our findings suggest a drastic change in the positioning of critical *CLV3* regulatory regions between Arabidopsis and tomato. Despite the finding that large 5’ deletions of *SlCLV3* alone recapitulate a null phenotype, 3’ mutations still have weak phenotypes, suggesting that 3’ CREs could interact with specific 5’ CREs. We tested this by combining 3’ mutations with smaller 5’ mutations isolated to specific regions. We previously found that deletions in two conserved regions 5’ of *SlCLV3* (designated R1 and R4), each have a weak effect on tomato locule number, and their combination enhanced locule number additively and synergistically, though these effects were still weak and varied with specific perturbations (34).

Given the strong synergistic interactions between *AtCLV3* 5’ and 3’ regions and the functional relevance of R1 and R4, we tested whether either R1 or R4 interacted with regions 3’ of *SlCLV3*. We first performed sequential CRISPR-Cas9 targeting of the *SlCLV3* 3’ region in the background of an R4 deletion (R4-5) (**Fig 5A**). From 10 alleles, six carried a range of small 3’ deletions and had little or no effect on locule number compared to the R4-5 deletion allele alone. The remaining four alleles had larger deletions affecting multiple 3’ CNSs and weakly enhanced locule number compared to R4-5 (R4-5 + Reg3’*^g^*, R4-5 + Reg3’*^h^*, R4-5 + Reg3’*^i^*, and R4-5 + Reg3’*^j^*) (**Fig 5A**).

**Fig 5.**
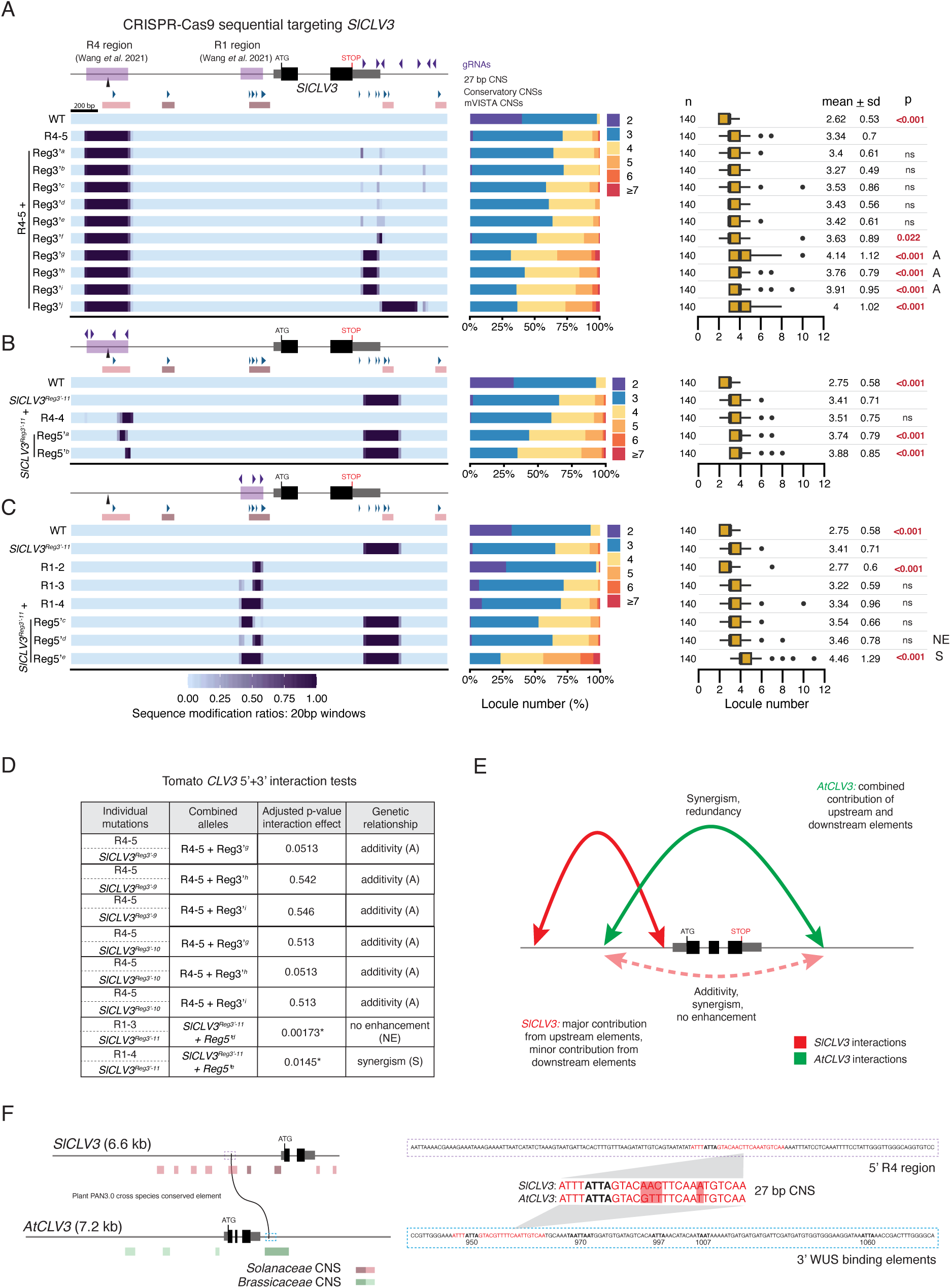
Combined mutagenesis of regions upstream and downstream of *SlCLV3* confirm the primary role of the upstream region in *SlCLV3* regulation. (A) Heatmap representation of alleles generated from sequential CRISPR-Cas9 targeting with the *SlCLV3* 3’-gRNA array, in the background of the fixed R4-5 mutant. Locule number quantifications are represented by stacked bar plots and box plots. (B) Heatmap representation of alleles generated from sequential CRISPR-Cas9 targeting with the *SlCLV3* R4-gRNA array, in the background of the fixed *SlCLV3^Reg3’-11^* allele. Locule number quantifications are represented by stacked bar plots and box plots. (C) Heatmap representation of alleles generated from sequential CRISPR-Cas9 targeting with the *SlCLV3* R1-gRNA array, in the background of the fixed *SlCLV3^Reg3’-11^* allele. Locule number quantifications are represented by stacked bar plots and box plots. (D) Interaction tests performed between combined 5’+3’ alleles and similar individual 5’ and 3’ mutants. p-values of the interaction effect were adjusted for multiple comparisons. (E) Model summarizing the relative contribution of the 5’ and 3’ region, as well as their interactions, to the regulation of *SlCLV3* and *AtCLV3.* (F) A conserved 27 bp sequence which overlaps with the distal R4 region in the tomato 5’ (outlined by a purple dashed box), and a known WUS TFBS in the Arabidopsis 3’ (outlined by a blue dashed box). The DNA sequences within these regions are shown, with the 27 bp sequence in red, nucleotide mismatches highlighted in red, and the core ATTA WUS binding element in black. All five of the previously characterized *AtCLV3* 3’ WUS binding elements are also bolded and named according to their position, as defined previously (35). (A) – (C) Box plots show the 25^th^, 50^th^ (median) and 75^th^ percentiles, with outliers as black points. Number of fruits sampled (n) is shown to the left, and mean and standard deviation (sd) are shown to the right. The R4 and R1 regions previously defined are highlighted by purple boxes on the *SlCLV3* 5’ non-coding sequence (34). A, additive interaction type. NE, no enhancement. S, synergistic interaction type. Purple arrowheads, gRNAs. Black arrow, 27 bp conserved element. Blue arrowheads, Conservatory CNSs. Red bars, mVISTA CNSs. Two-sided Dunnett’s compare with control tests were performed to compare WT and sequentially edited alleles to (A) R4-5, (B) *SlCLV3^Reg3’-11^*, or (C) *SlCLV3^Reg3’-11^*.

In the reciprocal experiment, we targeted the R4 or R1 region in the background of the weak 3’ allele *SlCLV3^Reg3’-11^*. All combined alleles targeting R4 were weakly enhanced by less than one locule compared to the effects of individual mutations, though sequential mutations in the R4 region were different than the R4-4 allele, preventing an accurate evaluation of interactions (**Fig 5B**). Combined R1 and 3’ mutations had various interactions (**Fig 5C, S2 Fig**). The 3’ deletion allele combined with partial deletions in the R1 region (allele *SlCLV3^Reg3’-11^* + Reg5’*^d^*) did not show an enhancement in locule number compared to *SlCLV3^Reg3’-11^* (**Fig 5D, S2 Fig**). However, the 3’ deletion allele combined with a full deletion of the R1 region overlapping all CNSs exhibited the largest increase in locule number of all combined alleles, but this effect was again weak (allele *SlCLV3^Reg3’-11^* + Reg5’*^e^*) (**Fig 5D, S2 Fig**). Statistical analyses confirmed additive effects except for *SlCLV3^Reg3’-11^* + Reg5’*^e^*, which was weakly synergistic (**Fig 5D, S2 Fig**). Thus, in contrast to our findings in Arabidopsis, combined mutations in 5’ and 3’ regions of tomato *CLV3* are predominantly weak in their enhancement of locule number and additive in effect, demonstrating most *cis*-regulatory function in tomato is restricted to upstream CREs.

Our data show that while conserved regions are present both upstream and downstream of *AtCLV3* and *SlCLV3* their phenotypic relevance and interactions are lineage-specific (**Fig 5E**). Despite an overall lack of non-coding sequence alignment between distantly related orthologs, short 10-30 bp sequences of similarity can often still be discovered, albeit in drastically altered arrangements (10,11,24). To identify such short conserved sequences between upstream and downstream regions of tomato and Arabidopsis *CLV3*, including the possibility that conservation might exist upstream in one species and downstream in the other, we performed a cross-species analysis between the entire 5’ and 3’ regions of *SlCLV3* and *AtCLV3* using PlantPan3.0 (40). This comparison exposed a nearly identical 27 bp sequence found in the distal upstream R4 region of *SlCLV3* and the proximal downstream region of *AtCLV3* (**Fig 5F**). Notably, this sequence also contained a WUS binding site; the ATTA motif to which WUS is known to weakly bind is completely conserved within the 27 bp sequence (35). Importantly, alleles that remove this sequence individually or in combination with other upstream or downstream mutations consistently showed phenotypic effects, and it’s possible that other TFBSs are also contributing beyond the WUS site (**Figs 2C, 3B, 3C, 3D, 4A, 5A**).

## Discussion

Although there are cases of remarkably deep conservation of non-coding sequence (41,42), divergence in the *cis*-regulatory sequence of otherwise conserved genes is a more common and widespread characteristic of distantly related orthologs (18). Despite *cis*-regulatory sequence divergence, transcription factors, their DNA binding domains, and their sequence preferences are commonly highly conserved (18). Thus, a likely explanation for the prevalence of *cis*-regulatory sequence divergence is that CREs can maintain transcription factor binding for various arrangements of TFBSs, allowing drastic shifts in overall sequence organization and grammar without compromising function. *AtCLV3* is regulated by WUS transcription factor binding, and multiple WUS motifs can be found in *SlCLV3* 5’ and 3’ regions, including within the 27 bp sequence conserved with Arabidopsis (**Fig 5F**) (35). Experiments such as ChIP-seq could establish whether the role of WUS in tomato *CLV3* regulation is direct, as in Arabidopsis, as well as the precise locations of functional WUS binding sites, which reflect only a subset of those found computationally and could also vary between species (35,43). Such data, including from other transcription factors, could shed light on the species-specific redistribution of *CLV3* CREs, which may contain WUS binding sites among other yet to be defined functional TFBSs that may be shared but differentially positioned and bound by their cognate TFs. In support, studies have shown that orthologous CREs can drive similar expression patterns in distantly related species despite extreme sequence divergence, likely through shared transcription factor motifs (10,11,24).

Another proposed constraint on CRE evolution is that although the binding sites of specific transcription factors are shuffled, their general genomic position (5’, 3’, intronic, exonic, within a UTR) relative to the gene may be conserved, thus preserving basic relative genetic and physical interactions among transcription factors and promoters (10,11,26,44). These studies suggest that modification of pre-existing CREs may be a more common evolutionary mechanism than CRE gains and losses. Contrary to this view, our results indicate that the genomic position of functional CREs contributing to the locule number phenotype were rearranged relative to the *CLV3* coding sequence between Arabidopsis and tomato during evolution. In the context of essentially no shared sequence identity, these rearrangements underscore that *CLV3* CREs lack even shared identity by functional synteny. Thus, while TFBS identity may often be constrained to ensure proper expression patterns, this does not necessitate the maintenance of 1:1 orthologous CREs.

Our results here and in past studies have demonstrated remarkable robustness of *cis*-regulatory regions to mutagenesis (24,34,37). This is a common feature of developmental processes, which are often canalized such that development is robust to genetic or environmental perturbations, such as the widespread and species-specific proliferation and hyper-variability of transposable element content in plant *cis*-regulatory regions (2). At least a portion of this robustness stems from the extensive modularity of *cis*-regulatory regions – namely, multiple CREs and their higher order, and often redundant, interactions in the control of gene regulation. We find extensive evidence of redundant interactions between CREs upstream and downstream of *AtCLV3*, and upstream of *SlCLV3* (**Fig 3**) (34). We observe a common trend where strong phenotypic effects often require mutations in multiple functional CREs. This trend extends beyond the *CLV3* locus, as large engineered perturbations to a 2.6 kb region upstream of tomato *WUS* also produced surprisingly weak phenotypes, indicating the existence of additional CREs beyond this region that function redundantly or in parallel in gene regulation (34). Redundancy can be encoded in CREs in various ways, including incorporating multiple binding sites for the same transcription factor within the same enhancer, and/or possessing multiple enhancers with the same function (i.e. redundant enhancers, also called shadow enhancers) (45). Evidence for both are prevalent in *Drosophila* development (12,13,46–48). Thus, multi-step mutations in multiple CREs may often be necessary during evolution to overcome dosage thresholds and elicit substantial phenotypic divergence for selection to act upon. However, it is also possible that we have engineered cryptic variants by perturbing CREs that buffer gene expression only in specific environmental conditions not yet assessed (49,50). Thus, robustness within *cis*-regulatory regions often allows sequence divergence to occur without sufficiently compromising expression to impact phenotype.

Engineering quantitative trait variation in crop species would benefit from a set of guiding principles, including reliable techniques to predict the most promising functional CREs and their relationships. The *cis*-regulatory code remains notoriously difficult to decipher; however, non-coding sequence conservation is considered a reasonable predictor of regulatory function (1). Our study underscores the limitations of CNS analyses, especially in the detection of CREs across very large distances. While functional TFBSs may often be conserved across deep time, their small size, variable weight of importance of specific residues, and altered sequence context can make them difficult to detect via traditional alignment methods (16). Although we show that the relative genomic positioning of functional CREs is not maintained between Arabidopsis and tomato *CLV3*, this re-organization is partially predictable based on in-family CNSs, which are evenly distributed between the 5’ and 3’ in Arabidopsis, and biased towards the 5’ in tomato. Furthermore, we found that perturbations of in-family CNSs were associated with mutations having phenotypic effects, but the magnitude and functional relevance of these effects was unpredictable and species-specific. Thus, while in-family conservation may be a useful tool to guide CRE engineering, the relevance of these regions can differ on a gene-by-gene basis, indicating the importance of considering the full repertoire of potential CREs in each species of interest when determining target sites for crop engineering.

## Materials and Methods

### Plant material, growth conditions and phenotyping

Seeds of *Solanum lycopersicum* cv. M82 from our own stocks were used as the background for WT and CRISPR-Cas9 tomato mutagenesis experiments. During initial allele isolation, tomato plants were sown and grown in 96-well flats for ∼4 weeks before being transplanted to pots, and grown in greenhouse conditions. The greenhouse operates under long days (16h light, 8h dark) with natural and artificial light (from high pressure sodium bulbs ∼250 umol/m^2^), at a temperature between 26-28°C (day) and 18-20°C (night), with relative humidity 40-60%. For phenotyping, tomato plants were sown and grown in 96-well flats before being transplanted to Uplands field at Cold Spring Harbor Laboratory. Plants in the field were grown under drip irrigation and standard fertilizer regimes. For each unique genotype, locule number was quantified from 140 fruits, taken from 7-12 individual plants. Seeds of *Arabidopsis thaliana* (ecotype Col-0) from our own stocks were used as the background for WT and CRISPR-Cas9 mutagenesis experiments. Arabidopsis plants were germinated on ½ MS plates and transplanted to 32-well flats for growth. During initial allele isolation, plants were grown in growth chambers under long days (16h light, 8h dark) at 22°C and light intensity ∼100 umol/m^2^. For phenotyping, Arabidopsis plants were grown on ½ MS plates in a growth chamber for 1 week (continuous light, 22°C, ∼100 umol/m^2^) before being transplanted to 32-well flats and grown in greenhouse conditions. The greenhouse for Arabidopsis growth operates under long days (16h light, 8h dark) with natural and artificial light, at a temperature between 20-25°C. For each unique genotype, locule number was quantified using the stereo microscope for 140 siliques, taken from 7-10 individual plants. Raw locule counts are available in **S1 Data.**

### CRISPR-Cas9 mutagenesis, plant transformation, and selection of mutant alleles

Generation of transgenic tomato with CRISPR-Cas9 mutagenesis was performed as previously described (51). Briefly, gRNAs were designed with Geneious Prime (https://www.geneious.com). The Golden Gate assembly method was used to clone gRNAs into a binary vector with Cas9 and kanamycin selection (37,52). Binary vectors were introduced into tomato plants through *Agrobacterium tumefaciens* mediated transformation in tissue culture (53). Transgenic plants were screened for mutations using PCR primers surrounding the gRNA target sites. PCR products were screened for obvious shifts in size by gel electrophoresis, and mutations were characterized by Sanger sequencing. First or second generation transgenics (T0 or T1) were backcrossed to WT to eliminate the Cas9 transgene and purge the genome of potential off-target mutations. F2 or F3 plants from these crosses that were homozygous for the CRISPR-induced mutation were used for phenotypic analysis. Generation of binary vectors for Arabidopsis CRISPR-Cas9 mutagenesis also utilized the Golden Gate assembly method. Arabidopsis constructs used an intronized Cas9 previously demonstrated to increase editing efficiency (54). The intron-Cas9 (L0 pAGM47523) was cloned with RPS5a promoter (L0 pICH41295) and NOS terminator sequence (L0 pICH41421) into the L1 plasmid pICH47822. This was assembled into the L2 vector pAGM4723 with NPTII for kanamycin resistance (pICSL70004 in L1 pICH47732), pFAST-R selection cassette (pICSL70008 in L1 pICH47742), and the gRNAs (each with U6 promoter and gRNA scaffold). Arabidopsis plants were transformed with binary vectors using *Agrobacterium tumefaciens* floral dip (55). Transgenic seed was selected by fluorescence, germinated on ½ MS plates, and transferred to soil at 7 days post germination, after which plants were subjected to a heat cycling regime that fluctuated between 37°C for 30 h and 22°C for 42 h over the course of 10 days. This protocol was previously described to increase Cas9 editing efficiency in Arabidopsis (56). Following heat treatment, flower DNA was genotyped by PCR for mutations in the target region, and individuals with evidence of editing were counter selected by fluorescence for absence of Cas9 and grown in the next generation for screening of stabilized mutations. T3 or T4 plants homozygous for the CRISPR-induced mutation were used for phenotypic analysis. All CRISPR-Cas9 alleles generated in this study are described in **S2 Data.** All gRNA and primer sequences are listed in **S2 Table.**

### *Cis*-regulatory sequence conservation analyses, TFBS prediction, and Plant PAN3.0 cross species analysis

Within-family conservation analysis was performed to predict conserved non-coding sequences within the 5’ and 3’ of *CLV3* in Arabidopsis and tomato that were shared among several Brassicaceae and Solanaceae species, respectively. The closest *CLV3* ortholog from each species was determined based on the ortholog with the greatest similarity to Arabidopsis or tomato *CLV3* within the 5’ and 3’ regions. Forty kb of sequence upstream and downstream of the *CLV3* ortholog was extracted, and aligned to Arabidopsis or tomato *CLV3* using mVISTA Shuffle-LAGAN (http://genome.lbl.gov/vista/mvista/submit.shtml) (33). Conservation was calculated in 20 bp windows, with an 85% similarity threshold. Conservatory CNSs were obtained from The Conservatory Project (www.conservatorycns.com) (24). TFBSs were predicted by scanning the Arabidopsis and tomato *CLV3* 5’ and 3’ regions for motifs using FIMO in the MEME suite (http://meme-suite.org/doc/fimo.html) (57). Position frequency matrices for known plant transcription factors were obtained from the JASPAR CORE PFMs of plants collection 2022 (58). A p-value cutoff of 0.00001 was used to predict TFBSs. For analysis of shared transcription factor motifs between Conservatory CNSs, Brassicaceae or Solanaceae family CNSs within the 5’ and 3’ of *CLV3* were stitched together, separated by 20 generic “N” residues, and scanned for motifs in FIMO, with p-value cutoff of 0.001. Motifs present within the family CNSs of both Arabidopsis and tomato were identified from this output. To search for short, conserved non-coding sequences shared between Arabidopsis and tomato *CLV3,* the Plant Promoter Analysis Navigator (PlantPAN) 3.0 cross species analysis function was used (http://PlantPAN.itps.ncku.edu.tw) (40). The Arabidopsis and tomato *CLV3* gene with 5’ and 3’ regions were used as input.

### Statistical methods

Pairwise comparisons between various alleles were performed using two-sided Dunnett’s compare with control tests. A p-value cutoff of <0.05 was used. For testing the genetic interaction between 5’ and 3’ mutations, a linear model was used. Each four-way comparison (between WT, single 5’ allele, single 3’ allele, and the combined 5’+3’ allele) was modelled with a linear model in R with interaction effect included (59). A p-value of <0.05 was used as a cutoff for a significant interaction effect. P-values were adjusted for multiple comparisons using the Benjamini-Hochberg method in R.

## Supporting information

**S1 Table. Division of conserved sequences and TFBSs upstream and downstream of *AtCLV3* and *SlCLV3*.**

**S2 Table. Oligos used in this study.** gRNAs used in Arabidopsis CRISPR, gRNAs used in tomato CRISPR, and genotyping/sequencing primers.

**S1 Data. Arabidopsis and tomato locule phenotyping raw counts.**

**S2 Data. CRISPR-generated mutations in this study.**

## Acknowledgements

We thank members of the Lippman laboratory for comments, discussions, and assistance with phenotyping; J. Van Eck and Q. Jiang for performing tomato transformations; and T. Mulligan, K. Schlecht, A. Krainer, B. Fitzgerald and S. Qiao for assistance with plant care. This research was supported by the Cold Spring Harbor Laboratory School of Biological Sciences, and the Natural Sciences and Engineering Research Council of Canada Postgraduate Scholarships – Doctoral program to D.C., the National Institute Of General Medical Sciences of the National Institutes of Health under Award Number K99GM149939 (the content is solely the responsibility of the authors and does not necessarily represent the official views of the National Institutes of Health) to S.Z., and the Howard Hughes Medical Institute and National Science Foundation Plant Genome Research Program grant IOS-2129189 to Z.B.L.

## Author contributions

D.C. conducted the experiments, prepared the figures and wrote the manuscript. S.Z. developed the Arabidopsis CRISPR-Cas9 workflow methodology. Z.B.L. conceived and supervised the research, and wrote the manuscript. All authors read, edited, and approved the manuscript.

## Competing interests

Z.B.L. is a consultant for and a member of the Scientific Strategy Board of Inari Agriculture.

## Data availability

All data are available in the main text, supplementary materials, and source data.

---

**Fig S1.**
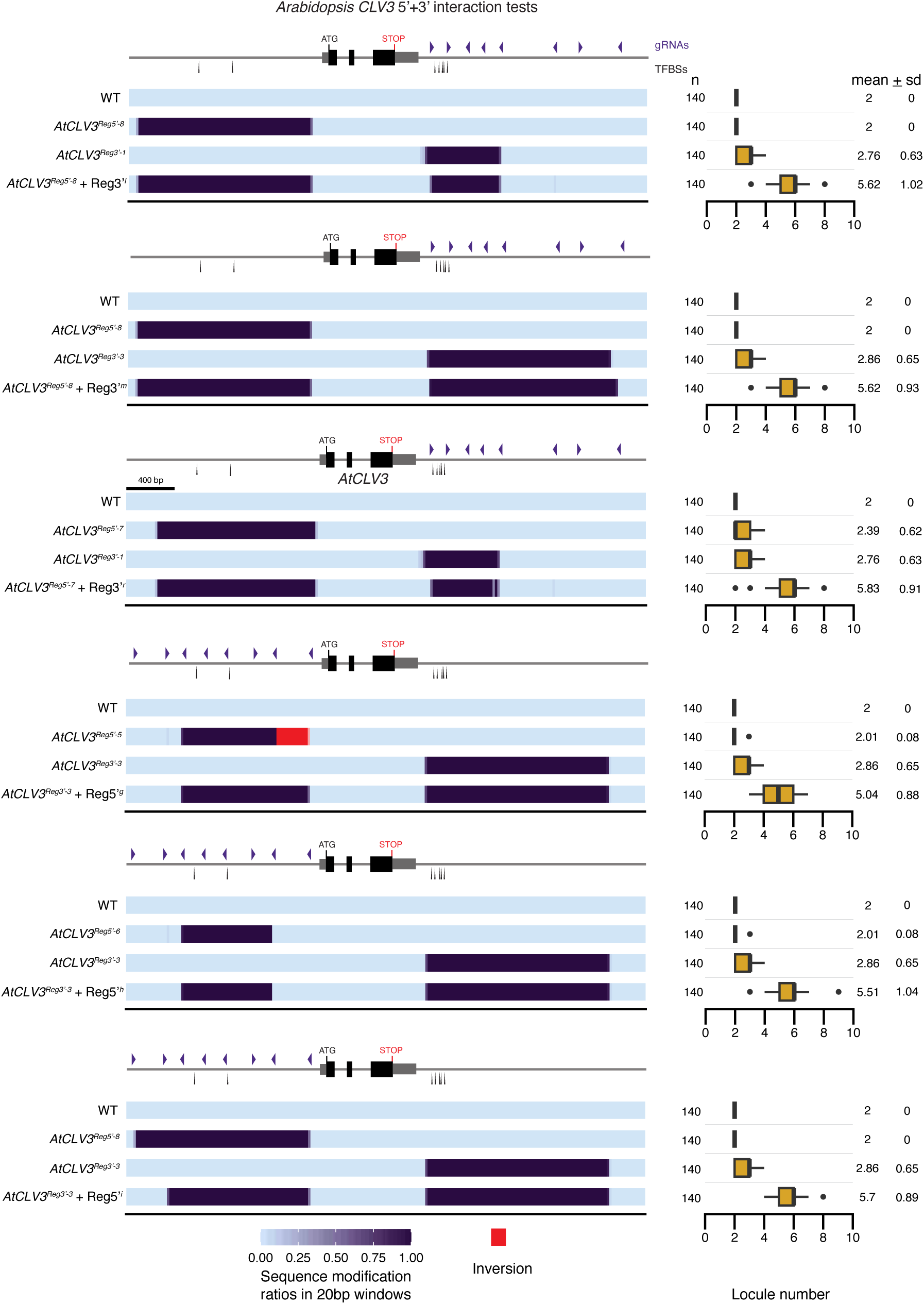
Arabidopsis *CLV3* alleles used for interaction tests. Heatmap representations of the *AtCLV3* alleles used in interaction tests, with their locule number quantifications. Each interaction test consisted of a linear model generated from the relationship among four alleles: one 5’+3’ combinatorial allele, one 5’ allele, one 3’ allele, and WT. Purple arrowheads, gRNAs. Black arrows, validated WUS and STM TFBSs.

**Fig S2.**
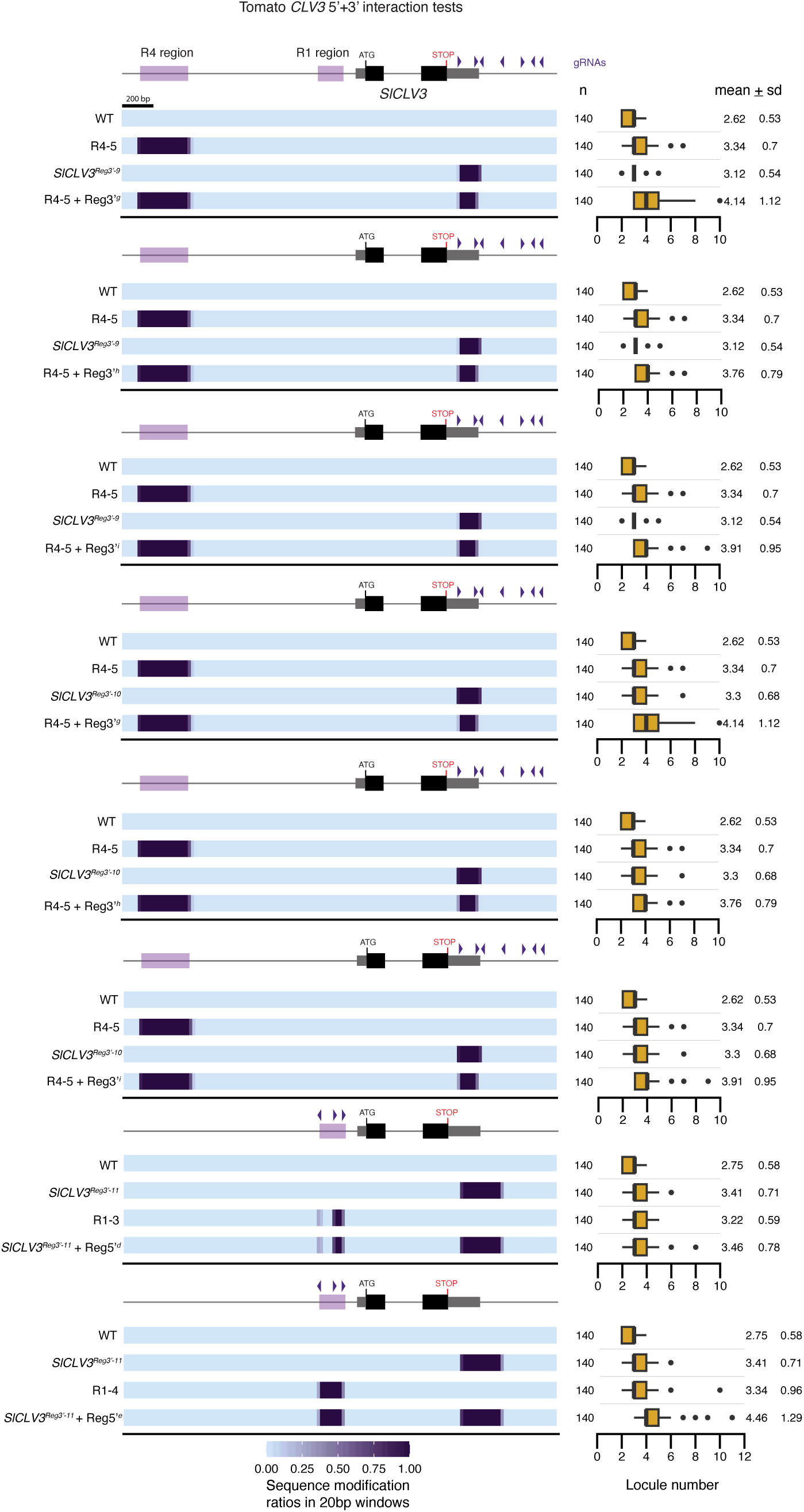
Tomato *CLV3* alleles used for interaction tests. Heatmap representations of the *SlCLV3* alleles used in interaction tests, with their locule number quantifications. Each interaction test consisted of a linear model generated from the relationship among four alleles: one 5’+3’ combinatorial allele, one 5’ allele, one 3’ allele, and WT. The R4 and R1 regions previously defined are highlighted by purple boxes on the *SlCLV3* 5’ non-coding sequence (34). Purple arrowheads, gRNAs.

